# Environmental DNA from ethanol eluent of flowers reveals a widespread diversity in cowpea associated animal communities in Hainan Island

**DOI:** 10.1101/2024.02.01.578371

**Authors:** Qi Chen, Huai-Liang Yu, Jun-Xian Lv, Xing Wang, Jin Li, Ming-Yue Wu, Cai-Hua Shi, Wen Xie, Xiang-Yi Kong, Guo-Hua Huang

**Author notes:** **Correspondence:** Xing Wang, Qiongtai Normal University, Haikou 571100,Hainan, China.; Guo-Hua Huang, Hunan Agricultural University, Changsha 410128, Hunan, China.

## Abstract

Cowpea (*Vigna unguiculata* (L.) Walp.), as an economical crop, is one of the important pillar industries of rural revitalization strategy in China. However, cowpea planting in China is often infested and damaged by many insects during growth, especially in Hainan region with a warm and wet tropical climate. Traditional monitoring methods with technical limitation could only detect a few common significant agricultural pests, how many kinds of species associated with cowpea is unknown. Here, we employed environmental DNA (eDNA) metabarcoding to characterize cowpea associated animal community-level diversity among six planting areas in Hainan. In all, 62 species were detected, of which 99.05% was Arthropoda, suggesting that Arthropods are the main groups interacting with cowpea. Moreover, we also detected 28 pests on cowpea, predominantly belonging to Thysanoptera, Lepidoptera, Diptera and Hemiptera, of which 20 pests were first reported and need more extra attention. Furthermore, clustering results indicated that there is a certain diversity of cowpea associated animals in different regions of Hainan, but the species composition was similar in the large planting areas due to the indiscriminate use of pesticides, which need further develop scientific pesticide applications to ensure adequate species diversity. This study represents the first molecular approach to investigate the cowpea associated animal communities and provides basic information for further scientific pesticide applications.

## Introduction

Biological monitoring is an essential condition for characterizing species diversity (van der Heyde et al., 2020, Vivien et al., 2015), assessing the ecological status of ecosystems (Andreasen et al., 2001, Vivien et al., 2015), and detecting the presence of invasive or pest species (Guareschi et al., 2020, Hardulak et al., 2020). As one of the most crucial terrestrial ecosystems on the planet, agricultural ecosystems have long been the most important resources for people to meet food, fiber and fuel needs (Swinton et al., 2007). Species diversity is a critical factor for the stability and sustainable development of agroecosystems (Ratnadass et al., 2011), thereby the biodiversity monitoring for agroecosystems is a long-term task in ecology-related fields. In agroecosystems, animals widely exist on nearly all continents as the largest quantities group of macroscopic organisms, functioning as biocontrol agents, pollinators and prey (Giribet & Edgecombe, 2012), playing a central role in the study of speciation, community ecology, biogeography and climate change (Pollard & Yates, 1994). Traditional animal community monitoring has largely been done with visual surveys or by passive sampling capturing specimens to identify and count species (Ji et al., 2022). Monitoring efforts requires researchers possessing rich knowledge on taxonomy and conducting careful analysis of morphological characteristics of specimens (Packer et al., 2009). However, these traditional survey methods are increasingly unable to meet the needs of large-scale diversity surveys. The reason is that different trapping methods often capture only a subset of actual biodiversity (Svenningsen et al., 2021, Krehenwinkel et al., 2022a) and large amounts of specimens (e.g., specimens of bees) could also make a biomonitoring extremely time-consuming (Gueuning et al., 2019).

As an important element of biomonitoring, environmental DNA (eDNA) is revolutionizing the way researchers monitor the biome inferring species presence and absence indirectly through molecular approaches (Coissac et al., 2012, Deiner et al., 2017, Valentini et al., 2016, Veilleux et al., 2021), whose surveys do not rely on visual observations, thereby simplifying species detection within inaccessible environments (Fediajevaite et al., 2021) and could identify simultaneously the large sets of taxa present in a bulk environmental sample (Taberlet et al., 2012b). To date, it has been turned out to be a reliable, reproducible, and time-effective tool in species discovery of the environmental samples from soil, scats (faeces), plant material, water, or air (Deiner et al., 2017, Taberlet et al., 2012a), which has been recommended as the primary approach for large-scale biomonitoring in plant species diversity of biodiversity coldspots (Liu et al., 2023), soil arthropod communities (Oliverio et al., 2018, Ustinova et al., 2021), fish diversity across river-lake connected system (He et al., 2022, Zhang et al., 2022), plants-pollinators interactions (Bell et al., 2022), and planktonic and benthic diatoms communities (Wang et al., 2019, Zhang et al., 2023). However, in agriculture, such superior technology is not widely used for plant-animal associations biomonitoring (Kestel et al., 2022), especially for that of the economic crops.

Cowpea (*Vigna unguiculata* (L.) Walp.) is an annual herbaceous plant belonging to the genus *Vigna* of the family Fabaceae (Wu & Mats, 2010), which widely distributed in Africa, Asia and Latin America (Ehlers & Hall, 1997). Cowpea is rich in nutrients, including vitamins, minerals (Ca, P, Fe), folate, thiamin and riboflavin, is a very important food source supplementing carbohydrates and proteins for people in developing countries (Mucheroa et al., 2009, Behura et al., 2015). In China, cowpea is widely cultivated over the country with planting area more than 0.67 million hectares and annual production for about 1.5 million tons, is an economically important crop in Fujian, Guangdong, Guangxi, Hainan, and Yunnan province where tropical or subtropical climate (Huan et al., 2015, Huan et al., 2016). However, cowpea in China is often infested and damaged by many insects during growth, especially in Hainan province with a warm and wet tropical climate (Huan et al., 2016, Wu et al., 2023). Moreover, with the indiscriminate use of pesticides, some small insects occurred are becoming more serious in recent years in Hainan region (Wu et al., 2023, Guo et al., 2023). Accurate species monitoring is a fundamental condition for precise pesticide application, which could avoid damage to some beneficial species (e.g. natural enemy) (Schmidt-Jeffris, 2023). Previous studies showed that *Megalurothrips usitatus* (Thysanoptera), *Maruca testulalis* (Lepidoptera), *Liriomyza sativae* (Diptera), and *Aphis craccivora* (Hemiptera) are main insects associated with cowpea occurred seriously in Hainan region (Li et al., 2022, Wu et al., 2023). However, these studies only focused on the pests, for other related species, how many kinds of that is unknown. Furthermore, we observed that there were at least nine species of insects potentially associated with cowpeas during our sample collection (Fig. 1). In view of this, we have reason to believe that the species related to cowpeas may be a large community which most of it has yet to be revealed on account of the limitation of monitoring way. Hence, it is necessary adopting a powerful approach to strengthen the monitoring of animal diversity associated with cowpeas and to accurately assess changes in biodiversity.

**Figure 1.**
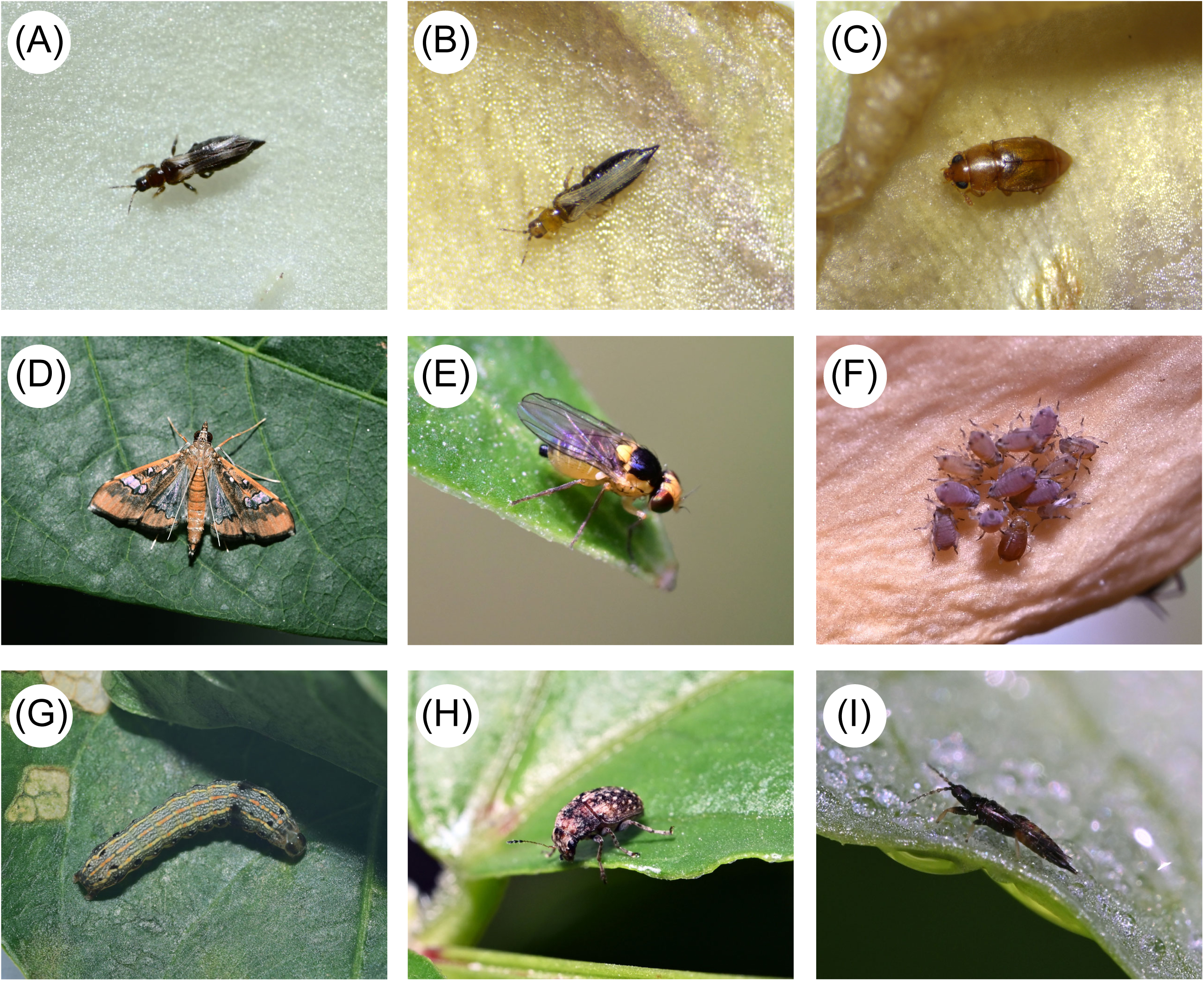
Photos of invertebrates associated with Cowpea found during our sampling. A representative for each species is shown. (A) *Megalurothrips usitatus* (Thysanoptera); (B) *Frankliniella intonsa* (Thysanoptera); (C) *Epuraea picinus* (Coleoptera); (D) *Maruca testulalis* (Lepidoptera); (E) *Liriomyza sativae* (Diptera); (F) *Aphis craccivora* (Homoptera); (G) *Spodoptera litura* (Lepidoptera); (H) *Araecerus fasciculatus* (Coleoptera); (I) *Haplothips* sp. (Thysanoptera).

Here, we focus on the planting areas of cowpea in Hainan province, then collect cowpea flowers gaining eDNA for animal community monitoring using DNA metabarcoding. This study aims to address the following questions: (i) how many other species are associated with cowpeas except for the major known pests? (ii) if species richness, whether there are differences in species composition between different planting regions. The present study would broaden the horizon of species diversity for cowpea associated animal communities and provide basic information for further application of precise pesticide in cowpea.

## Materials and methods

### Sample collection and DNA isolation

The study was carried out in the Hainan region (N 18°10’∼20°18’, E 108°37′∼111°05′) from February to August, 2023. A total of 46 samples of cowpea flowers were collected on altitude from low to high of cowpea growing area of Hainan province in the present study (Fig. 2, Table S1). For each sample site, a minimum of 50 cowpea flowers containing insects were collected and placed into a ziplock bag as a single sample, then added ∼ 200 ml of absolute ethanol to wash the flowers (Fig. 3A-3B). Insect specimens was recollected and used to carry out traditional morphological taxonomy while the rest ethanol liquid was filtered by a filtration device with 0.2 μm polytetrafluoroethylene (PTFE) microporous membrane. Environmental DNA was extracted from the microporous membrane following the protocol provided by SteadyPure Universal Genomic DNA Extraction Kit (Accurate Biotechnology Co. Ltd., Changsha, Hunan, China), with some modifications: increasing BuferLS-2 to 400 μl, proteinase K to 40 μl and RNase A to 20 μl was to enhance initial cell lysis and remove impurity (RNA and protein) in DNA extraction. DNA was preserved at -20 °C until PCR amplifications.

**Figure 2.**
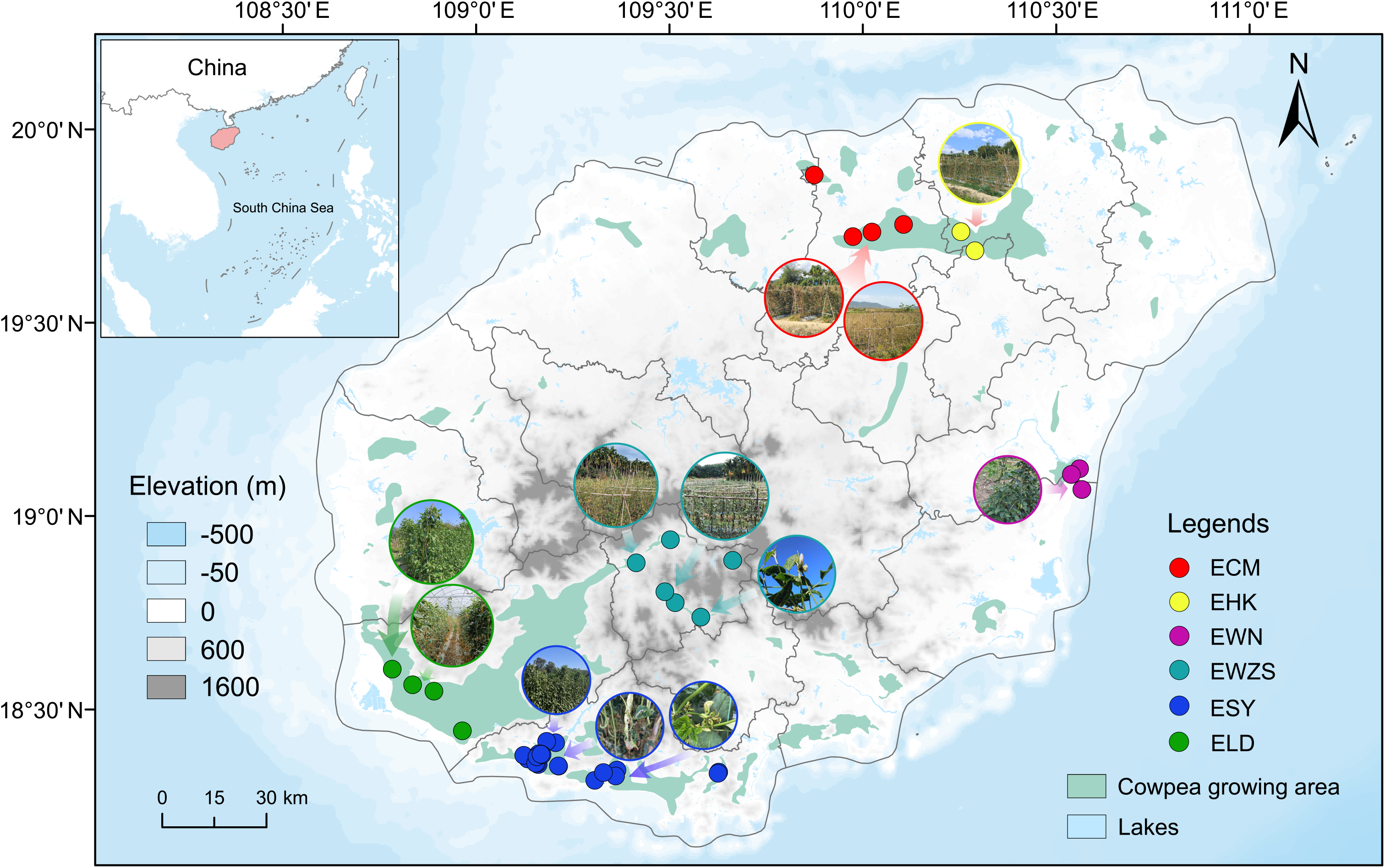
Map of Hainan region showing the sampling sites (N=46). Cowpea growing areas are marked based on our field visits and Google satellite map. Representative photos show the growth status of cowpeas at each sampling site.

**Figure 3.**
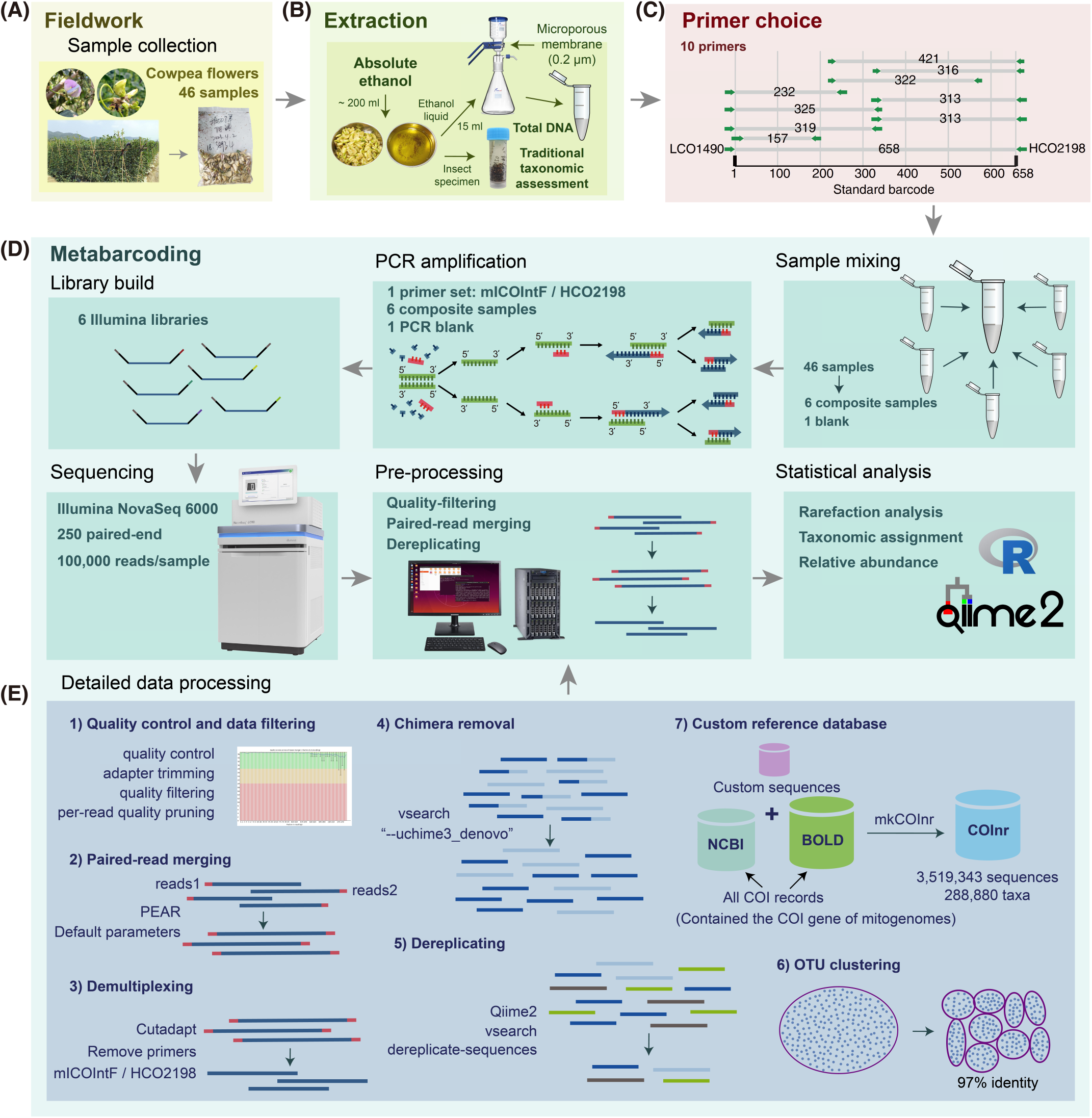
Study overview. (A) Samples collection from Cowpea growing area of Hainan. (B) eDNA sample and insect specimen collection. Using absolute ethanol washs the flowers. Insect specimen is used to carry out traditional morphological taxonomy and ethanol liquid is used to extract environmental DNA. (C) Selection of potential COI primer sets for DNA metabarcoding. Primer pairs shown are typically used/suggested combinations from the literatures (References were showed in supplementary material Table S2). (D) Metabarcoding pipeline. The 46 samples are processed as six composite samples for PCR amplification, amplicon libraries constructing and sequencing. Then a common bioinformatic pipeline are used for data standardization and statistical analysis. (E) Detailed bioinformatic pipelines for data pro-processing.

### Primer choice, PCR amplification, and sequencing

The goal of DNA metabarcoding is simultaneously identified all taxa contained in single environmental samples (Coissac et al., 2012, Taberlet et al., 2012b). This requires that the barcoding primer pair with suitable length for high throughput sequencing is universal and can be amplified the marker sequence of all taxa in samples for bioassessment (Elbrecht & Leese, 2017). Several COI barcoding primers with different levels of base degeneracy have been developed of which many are now used or could be suitable for animal community monitoring (Folmer et al., 1994, Geller et al., 2013, Iftikhar et al., 2016, Alberdi et al., 2017, Elbrecht & Leese, 2017, Kuntke et al., 2020, Xu et al., 2022). We carefully selected 10 COI primers from these literatures for primer evaluation by using PCR amplifications (Fig. 3C, Fig. S1, Table S2). Only the primer mICOIntF/HCO2198 (Beijing Tsingke Biotechnology Co., Ltd., Haikou, Hainan, China) showed good amplification bands, thus, which was selected as the primary primer in hereafter biomonitoring (Fig. S1). Amplifications were carried out in duplicates with a final volume of 50 μl. Each reaction contained 2 μl of the forward and reverse primer, 25 μl of 2×Taq PCR Mix (TIANGEN Biotechnology Co., Ltd., Beijing, China), 5 μl dd H_2_O (TIANGEN Biotechnology Co., Ltd., Beijing, China) and 16 μl of template DNA. Cycling conditions were as follows: initial denaturation at 94 °C for 3 min, 35 cycles of 94 °C for 30 s, 58 °C for 30 s, 72 °C for 1 min and final elongation at 72 °C for 5 min. The expected PCR product size (∼ 313 bp) was visualized on a 2% agarose gel. Then the amplified products were mixed for sequencing using Illumina NovaSeq 6000 platform with pair-end 250 bp (PE250) mode at Shanghai Biozeron Biological Technology Co. Ltd. (Shanghai, China).

### Construction of custom reference database

Reliability and accuracy of DNA based taxonomic assignment often relies on the wide taxonomic coverage of reference databases (Iwasaki et al., 2013). The NCBI (https://www.ncbi.nlm.nih.gov/) and BOLD (http://www.boldsystems.org/) are mainly two nucleotide databases used for animal taxonomic assignments in DNA metabarcoding. However, the data from these two major databases cannot be shared in real time. Many researchers, by contrast, prefer to submit their new marker sequences of known or new species to BOLD database, but some do not. Therefore, it is essential to integrate these two primary databases creating a custom database for more accurate taxonomic assignment of metabarcoding sequences. mkCOInr v.0.2.0 (Meglecz, 2023) was employed to create such custom database following operation document. The COI sequences were downloaded from these two nucleotide databases in November 13, 2023, respectively. Then using a series of semi-automated pipelines to format these COI sequences generating COInr database: a) retain CDS with gene and protein names corresponding to COI as well as the length range of 100-2000 nucleotides, b) remove the genes with introns, c) check the Latin names and normalize some ambiguous Latin names of sequences, and d) eliminate sequences that are substring of another sequence of the same taxID. For those new sequences in BOLD not present in NCBI database, the pipelines could not recognize and list the correct taxonomic information. We further fixed these errors through a custom script. Moreover, we also supplemented some new sequencing data of our laboratory into this COInr database. Finally, our newly constructed COInr database was available with Qiime format in Zenodo (10.5281/zenodo.10453096), which included 288,880 taxa with 3,519,343 sequences.

### Bioinformatic analysis

Raw data was carried out quality control using FastQC v0.11.9 (https://www.bioinformatics.babraham.ac.uk/projects/fastqc/), then filtered using fastp v0.23.2 (Chen et al., 2018), with adapter sequences trimmed referring to the self-provided Illumina adapter sequence database, also leading and trailing bases with quality below 30 were removed for each read (Fig. 3D-3E). Paired-end reads of clean data were merged using PEAR v0.9.11 with the default parameters (Zhang et al., 2014). Primer sequences (mICOIntF/HCO2198) were trimmed from the merged reads with Cutadapt v4.4 (Martin, 2011). Chimeric sequences were detected and removed using uchime3_denovo implemented in VSEARCH v2.22.1 (Rognes et al., 2016). Subsequently, we used the plugin VSEARCH of Qiime2 v2023.5 pipeline (Bolyen et al., 2019) to dereplicate sequences, then cluster into molecular operational taxonomic units (mOTUs) at 97% similarity. We further added filtering step to retain mOTUs that occurred in at least one sample and more than 10 times to obtain the final feature table. The taxonomic assignment was performed in Qiime2 pipeline on the basis of sklearn-based taxonomy classifier using our newly constructed COInr database. Sequencing depth and recovered diversity per sample was investigated using rarefaction curves with R package vegan v. 2.6-4 (Oksanen et al., 2022). Beta-diversity analysis was conducted using Bray-Curtis pairwise dissimilarity matrix within vegan R package, and visualized with nonmetric multidimensional scaling (NMDS) plots. Hierarchical clustering was performed to compare the differences in species composition of six groups of samples, and visualized with a heatmap using R package ComplexHeatmap v2.15.4 (Gu, 2022).

## Results

### Metabarcoding data

From February to August, 2023, a total of 46 samples of cowpea flowers were collected, then mixed into six samples for sequencing in the present study (Table S1). In PCR amplifications, none readings were observed for blank control (Fig. S1), indicating that there was no significant contamination throughout the experiment and that the experimental results could reflect the actual amplification of the sample. Illumina sequencing yielded a total of 4,435,485 reads from six samples (Table S3). After quality filtering, merging, denoising and chimera removal, a total of 4,428,974 effective reads were retained used for downstream analysis. Clustering with 97% similarity produced a total of 1532 mOTUs, and the corresponding sequence number accounted for 51.38% of the effective sequence. Based on our newly constructed COInr database, we detected 221 taxa in total, of which 160 taxa could be accurate identified and the corresponding sequence number of identified taxa accounted for 95.03% of the effective sequence within taxon sequences. Rarefaction curves reached the stationary phase well before reaching the maximum number of reads, indicated that the sequencing depth was sufficient to recover most cowpea associated animals (Fig. S2A). The rank abundance curves reflected the evenness and abundance of mOTUs in the six samples both horizontally and vertically (Fig. S2B), and showing that the species composition in different regions of Hainan has some differences to a certain extent.

### Taxonomic composition of eDNA communities

Taxonomic assignment was carried out based on sklearn-based classifier in Qiime2 pipeline using our newly constructed COInr database, which represented a surprised result about cowpea associated animal communities (Fig. 4, Table S4). At the phylum level, Arthropoda (99.05%) was the most represented phyla estimated by eDNA metabarcoding, while Nematoda (0.28%), Rotifera (0.57%), Mollusca (0.003%), and Chordata (0.001%) were also detected in small numbers, only 0.08% was unclassified (Fig. 4A, Table S4). At the class level, undoubtedly, Insecta (92.92%) was the most dominant class of cowpea associated animals, followed by Eurotatoria (0.57%), Hexanauplia (0.33%) and Chromadorea (0.2%), whereas 5.62% was unclassified (Table S4). At the order level, Thysanoptera (59.18%) and Coleoptera (23.63%) were two major orders identified, followed by Lepidoptera (6.14%), and Hemiptera (3.15%) (Fig. 4B, Table S4). At the family level, the dominating families were Thripidae (58.84%), Nitidulidae (20.58%), Crambidae (9.07%), and Reduviidae (2.44%), respectively, whereas 9.05% was unclassified (Table S4). At the genus level, a total of 33 genus were identified, of which *Megalurothrips* (52.69%), *Epuraea* (20.58%), *Frankliniella* (5.91%), *Maruca* (5.81%) and Rhynocoris (2.44%) was the main genus accounting for more than 87% of the relative abundance (Table S4). At the species level, *M. usitatus* (59.69%) and *Epuraea picinus* (20.58%) was the most represented species estimated by eDNA survey, followed by *Frankliniella cephalica* (5.75%), *Maruca* sp. (5.59%) and *Rhynocoris kumarii* (2.44%) (Fig. 4C, Table S4).

**Figure 4.**
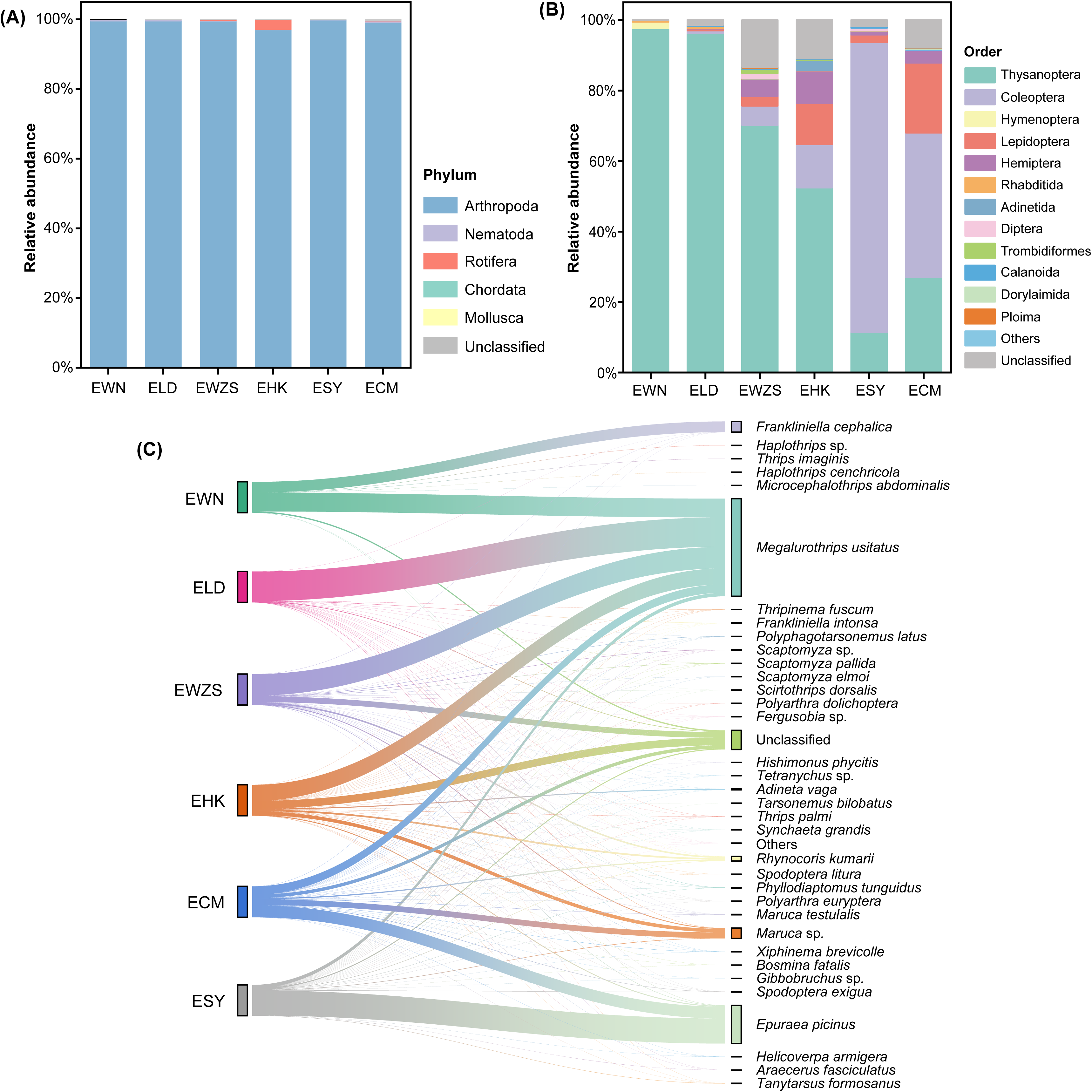
Relative abundance of Cowpea associated animal communities at (A) Phylum and (B) Order. (C) Sankey based on assigned taxonomy showing the species diversity of Cowpea associated animal communities.

### Estimation of species diversity of cowpea pests

We observed a great diversity of cowpea pests by collecting eDNA from flowers for taxonomic assignment based on bioinformatic method. The results showed a lot obvious differences in pest species detected among the 6 groups (Table S5). EHK exhibited the largest variety of pests (22), followed by EWZS (18), ESY (17) and ECM (16), whereas EWN (7) had the least variety of that. Among the pests, unquestionably, *M. usitatus* was the most abundant pests in quantity, the next were *F. cephalica*, *Spodoptera exigua*, *M. testulalis* and *F. intonsa*, while *Haplothrips statices* was the least (Table S4, Table S5). Moreover, it is great interesting that eDNA monitoring also detected some pests that had not been presented by traditional methods, such as *Araecerus fasciculatus*, *Hishimonus phycitis*, *Helicoverpa armigera*, *Cadra cautella* and *F. cephalica*. Furthermore, some species with certain abundance had also been observed, such as *E. picinus*, *Phyllodiaptomus tunguidus* and *Tanytarsus formosanus*, but it is uncertain whether they are cowpea pests (Table S5).

### Differences in species composition between groups

Hierarchical clustering of observed taxonomic groups associated with cowpea revealed relationships between the occurrence and abundance of the individual taxa and the locations (Fig 5). Differences in community composition between the six groups were correlated with the regional distribution to some extent, and that this signal is driven by the presence or abundance of several taxa (Fig 2, Fig 5). For example, some species were detected only in specific groups (Table S6), such as *Henosepilachna vigintioctopunctata* in EHK, *Bemisia tabaci* in ECM, *Tyrophagus putrescentiae* in EWZS, and so on, clusters of these species in heatmap were observed, and made significant contributions for the difference between different groups. Moreover, the groups from large growing areas of cowpea tended to cluster into a cluster, while the group from small growing areas of that did not, which indicated that the animal species composition was similar in large growing areas of cowpea. By comparison, however, from the UPGMA (Bray-Curtis distance) tree, four groups (EHK, EWZS, ELD and EWN) clustered into a cluster while the remaining two (ESY and ECM) formed a group when considering the abundance of taxa (Fig S3A). These results were supported by nonmetric multidimensional scaling analysis (NMDS) to some extent (Fig. S3B), but the slight difference between the two methods was also present, which EWN was grouped separately in NMDS analysis.

**Figure 5.**
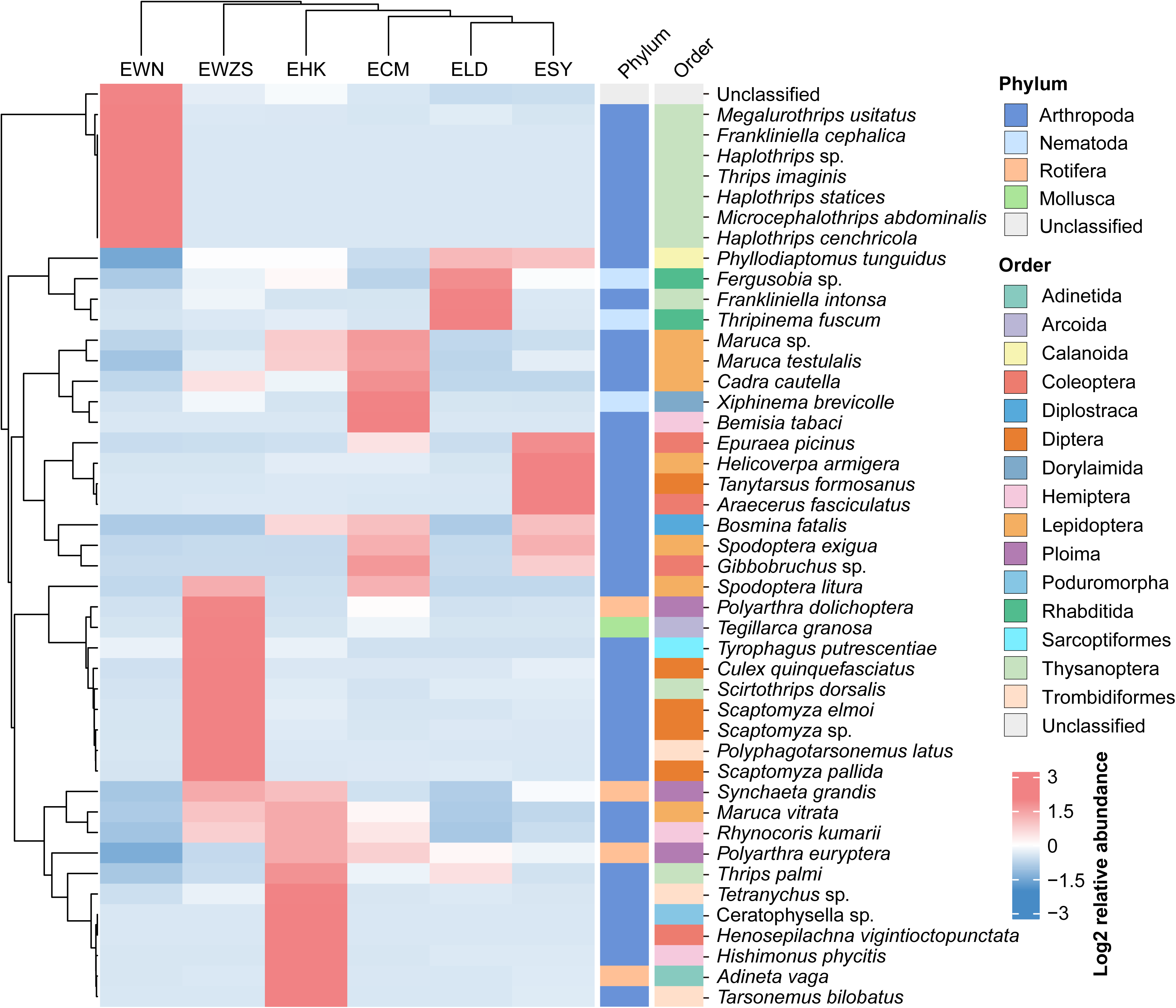
Heatmap exhibited different abundance of Cowpea associated species in different groups. OTU counts were converted to relative abundances, log-transformed and hierarchical clustering based on the average-linkage clustering methods in ComplexHeatmap.

## Discussion

The natural population densities of animals in agricultural ecosystems, especially arthropods, are associated with the variety and large quantities of cultivated crops (Thomas & Marshall, 1999, Andow, 1991). Only when the population density of one or more species reaches a certain scale will it attract the attention of agricultural practitioners (Wenda-Piesik & Piesik, 2020). Traditional monitoring for these species is mainly by setting traps (e.g. Malaise trap, light trap, flight interception trap, pitfall trap and so on) to capture a certain number of specimens, and then classify and count these specimens by morphological characteristics (Montgomery et al., 2021), which is tend to be lengthy and expensive, with the result that the timeliness and reliability of biomonitoring is greatly reduced. Recent development of eDNA metabarcoding just overcomes the limitations of the above traditional methods and has proved to be a feasible option for monitoring the animal communities with fast, standardized, and comprehensive features (Coissac et al., 2012, Taberlet et al., 2012a, Valentini et al., 2016, Deiner et al., 2017, Kestel et al., 2022). In the present study, a total of 46 samples were collected from the cowpea planting area of Hainan province for monitoring of cowpea associated animals using eDNA metabarcoding (Fig. 1, Table S1). The results showed a very abundant animal diversity (62 species, 37 of which can be accurately identified), which has greatly enriched our knowledge of cowpea associated animal communities (Fig. 4, Table S4). In the animal communities we detected, Arthropoda was the richest animals estimated by eDNA metabarcoding, revealed that the relationships between cowpea and animals is mainly focused on plant-arthropod interaction. Moreover, small numbers of other animal communities were also detected, such as Nematoda, Rotifera, and Mollusca (Fig. 4A, Table S4), but they may be the contamination whose eDNA came from the agricultural irrigation water. Furthermore, our eDNA survey revealed that Thysanoptera, Coleoptera and Lepidoptera were three major groups in the growing process of cowpeas (Fig. 4B, Table S4), exhibiting numerous overlaps with traditional methods (Li et al., 2022, Wu et al., 2023), which demonstrates the usefulness of eDNA metabarcoding for biodiversity characterization of cowpeas associated animals.

Information regarding the composition of pests is important for the ecological sustainable management and conservation biological control of crops (Gardarin et al., 2018, Hu et al., 2023). With the indiscriminate use of pesticides, the pests on cowpea planting in Hainan region gradually changed from large size to small size in morphology in recent years (Wu et al., 2023). Previous studies revealed that *M. usitatus*, *F. intonsa*, *Thrips palmi*, *Thrips hawaiiensis*, *L. sativae, Trialeurodes vaporariorum*, *M. testulalis*, *A. craccivora*, *Tetranychus cinnabarinus* and *B. tabaci* were the major pests on cowpea (Li et al., 2022, Wu et al., 2023). In this study, we detected a total of 28 pests on cowpea of Hainan using eDNA metabarcoding (Table S5). Although many of them (8 species) have been mentioned in previous studies (Li et al., 2022, Wu et al., 2023), most (20 species) are still being reported for the first time and need more extra attention, such as *A. fasciculatus*, *H. phycitis*, *H. armigera*, *C. cautella* and *F. cephalica* and so on. It is worth noting that *F. cephalica* is an exotic invasive species found in the plant *Bidens pilosa* in 2013 and mainly distributed in Guangdong, Guangxi and Hainan in China (Tong & Lv, 2013). Moreover, we found the population density of *E. picinus* (see Fig. 1C) was very high in the middle and late period of cowpea in Sanya (Fig. 4C, Table S4, Table S5). Such a high population density in one area is abnormal, and further increase monitoring efforts should be done, and do more in-depth research to re-examine whether this species has a risk of harming cowpeas.

The application of environmental DNA in ecology and conservation is to simultaneously detect the widest possible range of species from one or more habitats for characterizing past and present biodiversity patterns, and then exploring the mechanism and motivation of the phenomenon (Beng & Corlett, 2020). This requires researchers to accurately monitor species within a region, map their distributions, discover differences in species composition access habitats, and then design effective measures for this region for ecological management and conservation (Stockwell et al., 2003, Fisher & Owens, 2004). In the present study, differences in community composition between the six groups were observed and driven by the presence or abundance of several taxa (Fig. 5, Fig. S3), showed that there is a certain diversity of cowpea associated animals in different regions of Hainan. This may be correlated with local crop planting patterns and the amount of vegetation. However, the four groups from large growing areas of cowpea tended to cluster into a cluster, while the groups from small growing areas of that did not, suggesting similar species composition in these large planting regions, indicated that long-term cultivation of cowpeas, followed by the indiscriminate use of pesticides, could change the pattern of biological diversity. Therefore, under the long-term cultivation of cowpea, adopting effective ecological management, scientific pesticide application to maintain relatively rich species diversity, is still a crucial topic worth long-term discussion in cowpea industry.

This study based on environmental DNA metabarcoding has greatly expanded our knowledge of the diversity of cowpea associated animal communities, but it also has some shortcomings. One is insufficient sampling. We sampled only the flowers of the cowpea, thus some leaf pests (e.g. *L. sativae*) were not detected. A recent study based on plant-derived environmental DNA showed that there are significant differences of species composition of arthropod communities in different parts (e.g. flower, stem or root) of the same plant (Weber et al., 2023). In view of this, we believe that there is still a wider diversity of cowpea associated animal communities, which need for more extensive long-term monitoring of such communities in the future. The other is loss of complete growth cycle monitoring. Our samples were collected in the middle and late stages of cowpea growth (Fig. 2), and the results could not truly represent the animal composition in the early growth stages of cowpea. With the rapidly growing demand for large-scale biodiversity, method of metabarcoding is continuously developing and improving in recent years. Nascent surface eDNA-based survey methods provide a tractable mature solution, which could carry out a more convenient way for long-term monitoring of cowpea associated animals (Dangles & Casas, 2019, Krehenwinkel et al., 2022a, Krehenwinkel et al., 2022b).

## Data accessibility

Illumina sequence reads generated in this study have been deposited at China National Center for Bioinformation (CNCB) (https://ngdc.cncb.ac.cn/) under Bioproject xxx, and OTU table as well as custom reference database have been deposited in Zenodo (https://doi.org/10.5281/zenodo.10453096), respectively. The voucher specimens collected in this study are deposited at the Insect Museum of Hunan Agricultural University. Information on the samples can be found in supplementary material Table S1.

## Supporting Information

**Table S1.** Collecting information of samples used for environmental DNA surveys in the present study.

**Table S2.** 10 potential COI primer sets for primer evaluation using PCR amplifications for biomonitoring of Cowpea associated animal communities.

**Table S3.** Sequencing data statistics of biomonitoring of Cowpea associated animal communities.

**Table S4.** Relative abundance of all species of Cowpea associated animal communities.

**Table S5.** Arthropod species detection estimated by eDNA metabarcoding in this study.

**Table S6.** Unique species of each group detected by eDNA survey.

**Figure S1.** Evaluation of PCR amplifications. (A) primer evaluation; (B) samples evaluation.

**Figure S2.** Evaluation of sequencing results. (A) Rarefaction curve of mOTUs; (B) rank-abundance curve of mOTUs.

**Figure S3.** (A) Bray-Curtis dissimilarity dendrogram for the samples. (B) Nonmetric multidimensional scaling plots (NMDS) of Cowpea associated animal communities based on a Bray-Curtis pairwise dissimilarity matrix.

## Acknowledgments

The authors thank Mr. Ying-Qiu Li (South China Agricultural University, China) and Mr. Xiang Zou (Yangtze University, China) for their kind help in molecular biological experiment and preparation of experimental consumables. This work was supported by the National Key Research and Development Program of China (2022YFD1401200), and China Agriculture Research System (CARS-23-C08).

